# HSQC2STRUC: A Machine Learning Model for Protein Secondary Structure Prediction using Unassigned NMR Spectra

**DOI:** 10.1101/2023.10.09.561482

**Authors:** Jonas Dietrich, Peter Bellstedt

**Affiliations:** Institute for Inorganic and Analytical Chemistry, Friedrich Schiller University, Lessinstrasse 8, 07743, Jena, Germany; University of Zurich (UZH) & University Hospital Zurich (USZ), Institute of Clinical Chemistry, Rämistrasse 100, 8091, Zurich, Switzerland

## Abstract

Dynamic changes in the secondary structure content of proteins can provide valuable insights into protein function or dysfunction. Predicting these dynamic changes is still a significant challenge but is of paramount importance for basic research as well as drug development. Here, we present a machine learning-based model that predicts the secondary structure content of proteins based on their un assigned^1^H,^15^N-HSQC NMR spectra with an RMSE of 0.11 for *α*-helix, 0.08 for *β*-sheet and 0.12 for random coil content. Our model has been implemented into an easy-to-use and publicly available web service that estimates secondary structure content based on a provided peak list. Furthermore, a Python version is provided, ready to be integrated into Bruker’s TopSpin software or own scripts.

## 1. Introduction

Proteins play pivotal roles in natural processes, and their dysfunction is often linked to diseases including Parkinson’s and Alzheimer’s disease [28]. Developing appropriate therapies requires an in-depth understanding of the inextricable structure-function relationship of proteins. A major scientific breakthrough for the (in-silico) prediction of the three-dimensional structure of proteins was made by AlphaFold2 (AF) during the CASP14 challenge [11] and was followed by RoseTTAfold [4] and CollabFold [21] with similar approaches. Despite its remarkable success, AF has limitations arising from its training data and prediction process that prevent it from providing crucial insights into protein folding, stability and dynamics, rare folds, or ligand binding [14]. To obtain such essential insights, spectroscopic methods are still invaluable and provide information that often complements the results obtained from AF or similar approaches. Since determining a high-resolution 3D structure is often time-consuming, many scientific questions can be readily addressed by analyzing the (change) of secondary structure content. Although new optical methods are being actively developed [7; 15], gold-standard to estimate the amount of *α*-helical, *β*-sheet, random coil content in solution is circular dichroism (CD) spectroscopy [9; 19; 20; 33], whereas aggregated (or insoluble) protein structures are often characterized by infrared (IR) spectroscopy [34]. In addition, nuclear magnetic resonance (NMR) spectroscopy is widely used to characterize dynamic states of proteins [24], folding processes [2] and ligand binding [23]. The development of in-cell NMR provided a new perspective on structure-function relationships in their natural environment [29] and proved NMR as a versatile and complementary method for structural characterization. One of the most frequently recorded NMR spectra is the ^1^H,^15^N-HSQC [6], which is often referred to as a “fingerprint” spectrum since, despite its simplicity, it provides valuable information about the protein under investigation. However, the traditional NMR-based estimation of secondary structures (e.g., via TALOS+ [26]) requires ^15^N,^13^C double-labeled samples as well as the assignment of each peak to their respective amino acid. This renders the expression of protein expensive and the overall process time-consuming. First attempts to use *unassigned* ^1^H,^15^N-HSQC spectra to estimate secondary structure content served as a prove-of-concept [22]. Still, it was not competitive with other tools due to the limitation of a small data set of 64 proteins and the utilization of a simple linear regression model. Here, we present the development of a machine learning-based model trained on publicly available ^1^H,^15^N-HSQC spectra with experimentally determined protein structures from the Protein Data Bank (PDB) that estimates the secondary structure content of proteins based on their unassigned HSQC spectrum with reasonable accuracy.

## 2 Methods

### Data collection and filtering

All data originated from the PDB [5] or BMRB [10]. A subset of 8275 proteins with publicly available NMR data and with linked experimentally determined 3D structures were selected basedon a BMRB-to-PDB match list (retrieved from BMRB in November 2022) and was subsequently filtered by the following criteria: (1) Data availability: only entries with reported backbone ^1^H,^15^N chemical shift were considered. Only the first was considered if more than one BMRB entry was provided for a PDB structure. (2) Measurement conditions: Sample temperature during NMR experiments had to be between 273 K -310 K, sample pH 5-8, under- or overpressurized samples were excluded. If the conditions were not reported in an appropriate unit or not reported at all, the data were manually curated, and decisions were made on a case-by-case basis. (3) Sequence alignment: the *PairwiseAligner* (Biopython library) with an *open gap score* of -4 and an *extend gap score* of -0.01 was used to align the protein sequences provided by BMRB and PDB. Aligned sequences with less than ten amino acids were removed from the dataset. Additionally, the sequence alignment step removed all peaks originating from unaligned amino acids (resulting from e.g. additional purification tags). (4) Chemical shift range: only entries with ^15^N backbone chemical shifts between 90-140 ppm and ^1^H_*N*_ chemical shifts between 6-11 ppm remained in the dataset. (5) Type of biopolymer: DNA/RNA and DNA/RNA-protein complexes were excluded. The Biopolymer information was extracted from the PDB using the Python library pypdb. (6) Removal of selected proteins and binned duplicates: The binned spectra were compared using a distance cosine score function to avoid input data duplicates. The comparison was performed for a data set of 8x8 bins, resulting in a quadrant size of 6.25 ppm by 0.625 ppm for ^15^N and ^1^H, respectively. All spectral pairs with a cosine-distance-score of zero were identified and grouped based on their IDs. Then, all but one spectrum from each group was removed from the data set. The bin size 8x8 was chosen to ensure that all bins with a size of 8 or greater did not contain identical spectra. In addition, all *α*-synuclein and calbindin-related spectra (identified by using the respective search term on the BMRB web page) were removed from the dataset because the two prot eins were used as test cases.

### Data preprocessing

After the filtering step, the spectra were converted to a format suitable for machine learning and the secondary structure of the proteins was extracted as follows: Starting from the ^1^H,^15^N backbone spectra, all peaks between 90-140 ppm (50 ppm range for ^15^N) and 6-11 ppm (5 ppm range for ^1^H) were grouped (binned) into quadrants. The number of quadrants varied from 2 to 36 with a step size of 2 for both (^15^N and ^1^H) shifts resulting in 324 combinations that were tested during the model selection process. For the secondary structure determination, the PDB files were passed to the *DSSP module* [12] implemented in the PDB package of Biopython, which provided the Q3 secondary structure of every amino acid within the respective PDB entry. Output classified as polyproline structures (P) were added to the random coil fraction. Importantly, since the HSQC spectra often contained fewer peaks than expected from the full sequence given in the PDB entry, the percentage of helix, sheet, and random coil was calculated based on the aligned part of both sequences only and not on the full sequence.

### Model selection and optimization

The following 12 different machine learning models were tested during the model selection process: Bayesian Neural Network (BNN), CatBoost (CB), Decision Tree (DT), Elastic Net (EN), Extreme Gradient Boosting (XGB), Feed Forward Neural Network (FNN), Gradient Boost (GB), K-Nearest Neighbors (KNN), Light Gradient Boost Machine (LGBM), Random Forest (RF), Ridge Regression (RR) and Support Vector Machine (SVM). A simple Linear Regression (LR) model was used as a baseline for comparison. The initial hyperparameter settings for CatBoost and the two neural network architectures are given in Suppl. Listing 1-3. All other models used the default settings. Please note: Only the artificial neural network models and the CatBoost implementation can predict all three secondary structure states with one model, while all other models rely on separate models (one for each secondary structure state). Training and validation were performed using ten-fold cross-validation for all 324 binning combinations. Pearson and RMSE values were used as evaluation metrics (and are the mean values of the cross-validation runs). For each ML model, the best-performing binning sizes for the prediction of *α*-helix, *β*-sheet, and random coil content were selected independently and compared with each other. For the final CatBoost model, the input of the three best-performing binning sizes for the prediction of *α*-helix, *β*-sheet, and random coil content were combined into one, and the hyperparameters optimized using the optimization framework *Optuna* [1] with the TPEsampler used as the sampling algorithm (1000 runs per model). The independent models were optimized accordingly (without combining the input bin before).

## 3 Results

### Workflow

All remaining proteins in the dataset met the requirements to pass the upstream filtering process applied to the list of BMRB and PDB ID matches (Suppl. Fig. 1). The original list included 8257 proteins. Only proteins with a unique identifier and an ^15^N-HSQC spectrum available on the BMRB website were used (loss of 2438 proteins). Of the remaining 5181 proteins, only proteins measured at 1 ATM, pH 5-8, and temperature 273 K -310 K were included in the data set (loss of 1028 proteins). In addition, only proteins with peaks between 5 ppm to 11 ppm for hydrogen shifts and 90 ppm to 140 ppm for nitrogen shifts were further used (loss of 374 proteins). Based on a sequence alignment performed, proteins had to match more than 10 amino acids between the PDB sequence and the BMRB sequence of the protein (loss of 147 proteins). DNA and RNA biopolymers stored in the BMRB were also removed from the data set (loss of 92 biopolymers). The data set contained identically binned spectra that were grouped and all but one of the duplicates were removed from the data set (loss of 48 proteins). The selection of which proteins from the groups were included in the data set was random as the resulting secondary structure content should also be similar or the same. It was also ensured that all proteins used in the data set were unrelated to *calbindin*, and *α-synuclein* as these proteins were used as test cases afterward (loss of 5 proteins). Each of the different ML models (see Model selection) was trained and cross-validated on 90% (3581) of the remaining 3979 proteins, while the remaining 10% (398 proteins) were used as a test data set. The distribution of measurement conditions (temperature, pH), secondary structure abundance, and the NMR peak distribution of the (filtered) data set are shown in Suppl. Fig. 2, 3 and 4, respectively.

The cross-validation identified CatBoost as the best model for predicting all three secondary structure elements with a slight lead over LGBM and FNN (Suppl. Fig. 5). The scores and errors of all 12 ML models as well as the linear regression model are provided in Suppl. Tab. 1-3. The highest Pearson score (and lowest RMSE values) were obtained with the CatBoost model utilizing 14x10 bins (^15^N x ^1^H) for *α*-helix (Suppl. Fig 6), 10x8 for *β*-sheet (Suppl. Fig 7), and 16x12 for random coil (Suppl. Fig 8). Interestingly, different binning sizes were required to achieve the best predictions for each ML model and each secondary structure element (Suppl. Tab. 4).

### Optimization of the CatBoost model

Next, we checked if the prediction of the three secondary structure elements sum up to 100%. However, when using three independent CatBoost models (with their respective best-performing binning sizes), the cumulative secondary structure content varied between 70-130% (blue bars in Fig. 1). We therefore decided to combine the input of the three independent models into one single model, optimized the hyperparameters with Optuna (Supp. Table 5), and retrained this single model(s) with the optimized parameters. This combined final CatBoost model not only showed slightly better performance (Suppl. Tab. 6) in the 10-fold cross-validation but successfully learned to sum up all three secondary structure elements to 100% (orange bars in Fig. 1).

**Fig. 1.**
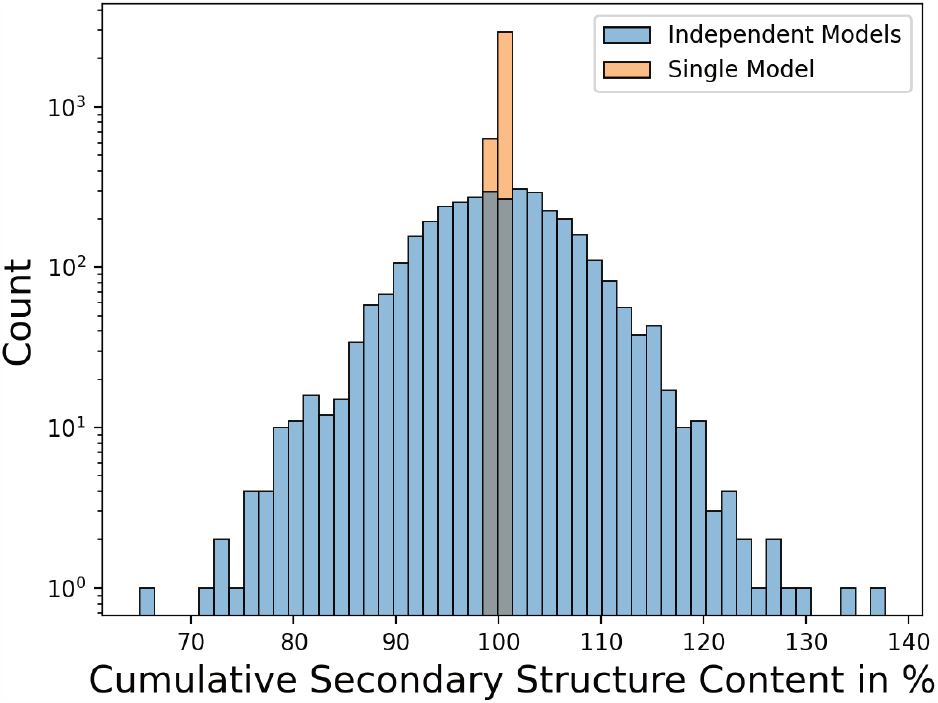
Cumulative secondary structure content prediction of the single ML-model combining three optimal binning sizes for each secondary structure element as input and three independent ML-models using only the optimal binning size for one secondary structure element each as input. Predictions are averaged from 10-fold cross-validation.

**Fig. 2.**
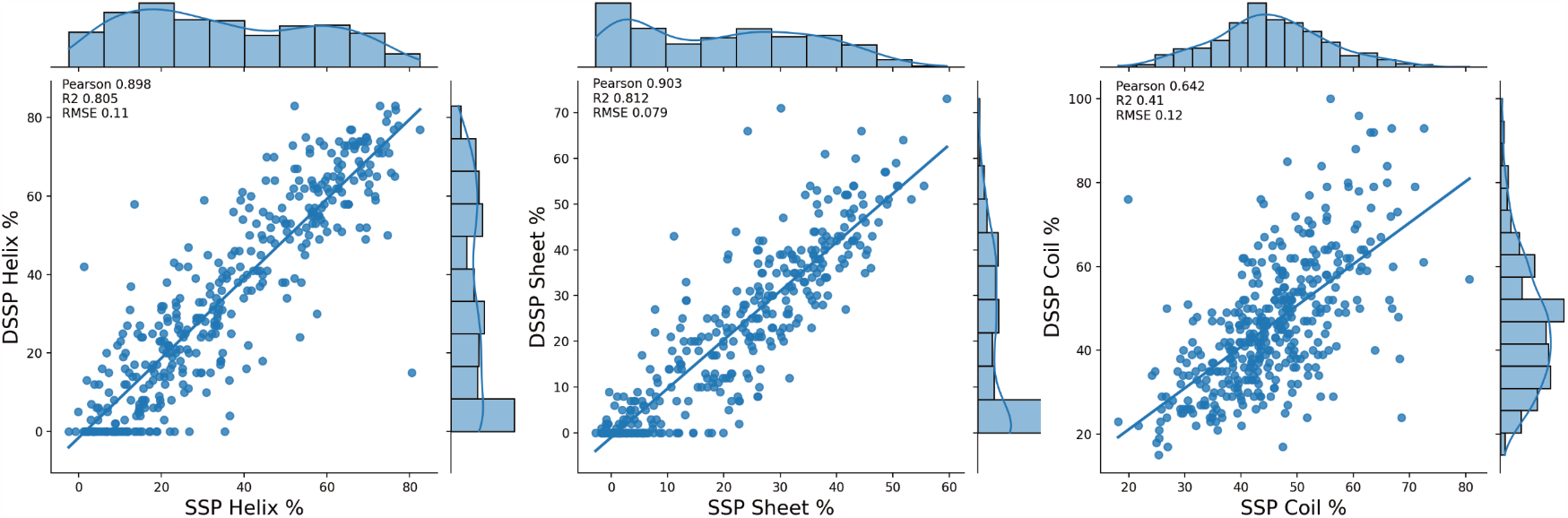
Performance of the final and optimized CatBoost model using the test data set. The “true” secondary structure content of each protein in the test data set extracted with DSSP is plotted against the secondary structure prediction (SSP) obtained with the model presented in this study. The Pearson score and Root Mean Square Error (RMSE) are given as measures to evaluate the model’s performance.

**Fig. 3.**
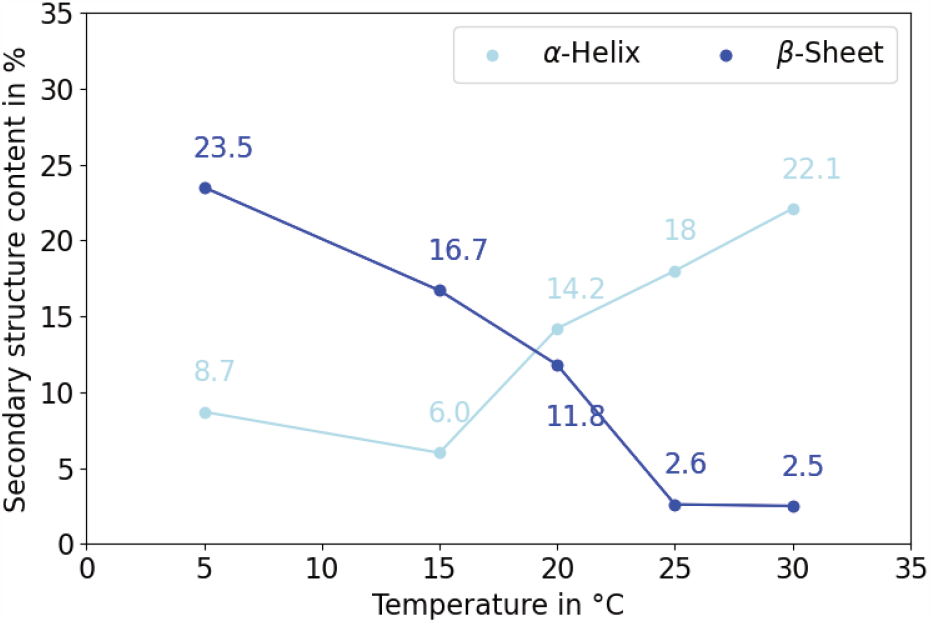
Predicted secondary structure content of *α* -synuclein measured at different temperatures

Next, we checked the performance of our final model using the test data set, which contained spectra/proteins that were never exposed to the model before. The optimized (single) CatBoost model demonstrated a high prediction performance, achieving a Pearson score of 0.90 for *β*-sheet followed by 0.89 for *α*-helix and 0.67 for random coil (Fig. 2). Although the predictions of our model only rely on a simple 2D NMR spectrum as input, the highest RMSE value here is 12%.

### Application

Our single CatBoost model is available for Python or can be readily integrated into the NMR analysis Software TopSpin (Bruker BioSpin). Peak picking is based on TopSpin’s peak-picking algorithm (pp command) or the manually selected peaks. Installation can be done by running the bash script available on GitHub in the TopSpin directory on the local computer (see Data Availability section for hyperlinks). Additionally, we integrated our model into an easy-to-use web service that accepts peak lists (* .csv) and provides a graphical output of the secondary structure content prediction as well as of the SHAP values [16] for each quadrant to understand the model’s decision. To allow for straightforward use, the input format corresponds to the peak lists generated using TopSpin’s export function. Additionally, backbone chemical shift files (*.csv) from the BMRB database can be uploaded without any further modification. Screenshots of the software integration as well as the web service, are provided in Supp. Figures 9 and 10, respectively.

## 4. Discussion

### Comparsion with previous work

The two most recent approaches in predicting secondary structure content with NMR data are TALOS+/ TALOS-N [26; 25] and PASSNMR [22]. TALOS is widely used in the scientific community, while PASSNMR is no longer publicly available but formed the basis for the here-developed model. PASSNMR was developed in 2006 and used unassigned ^15^N-HSQC and ^13^C-HSQC spectra to predict secondary structure content using a simple linear regression model. Our approach relies on ^15^N data to avoid the need for more expensive ^13^C-labeling, which would reduce its usability for users. Furthermore, the inclusion of ^13^C data would further decrease the already small data set that is available for training. PASSNMR used chemical shift ranges of 90 ppm to 140 ppm for nitrogen shifts and ppm to 11 ppm for hydrogen shifts, as is the case in this study. Furthermore, PASSNMR used a fixed binning size of 10x10. Here, improvements could be made by testing 324 different binning sizes from 2x2 to 36x36 with a step size of two. By using a much more extensive data set, a state-of-the-art machine learning model with optimized binning sizes, we could outperform the Pearson and RMSE values provided in the original PASSNMR publication. TALOS+ and TALOS-N, developed in 2009 and 2014, are hybrid systems for empirical prediction of protein *ϕ* and *ψ* backbone torsion angles and secondary structure content using a combination of six types of chemical shift assignments(HN, HA, CA, CB, CO, N) for a given residue sequence. TALOS-N is based on the same general artificial neural network (ANN) architecture as TALOS+, but the greater reliance on the ANN architecture results in higher prediction reliability. The TALOS architecture allows the use of amino acid sequences alone or in combination with assigned chemical shift values. Because our CB model relies only on unassigned ^15^N-HSQC spectra without any sequential information, the accuracy of both methods cannot be compared directly. The current commonly used model TALOS-N or TALOS+ for dynamic protein prediction requires high efforts and therefore reduces usability due to the inclusion of six different assigned chemical shift values needed for prediction. Our model is able to utilize low-cost ^15^N-HSQC spectra and allows the prediction of dynamic protein structure changes.

### Case studies

The overall goal of the study was to develop a tool to predict the secondary structure content of proteins using ^15^N-HSQC spectra. A key feature of NMR is its ability to monitor dynamic structural changes in proteins. To test the applicability of our model, we used NMR data of two key proteins involved in Parkinson’s and Alzheimer’s disease:calbindin and *α*-synuclein. Please note that our model was never trained/exposed to spectral data related to these two proteins to allow for a “real-world” -like case study (see Data collection and filtering).

#### Calbindin

Calbindin is a calcium-binding protein and is thought to play a critical role in protecting against cell death [13].

Five different entries were selected in the BMRB for calbindin, comprising two apo-forms as well as different ions bound. One of the two apo-forms also had a linked PDB entry and could therefore be calculated by DSSP and subsequently compared with the CB model predictions. The secondary structure predictions can be found in Table 1.

**Table 1.** Secondary structure content prediction for different calbindin-complexes.

| BMRB-ID | Form | Helix [%] | Sheet [%] | Coil [%] |
| --- | --- | --- | --- | --- |
| 19371 | Mg<sup>2+</sup> -bound | 69.6 | -0.4 | 30.8 |
| 19370 | Ca<sup>2+</sup> -bound | 56.9 | 6.5 | 36.6 |
| 6699 | La<sup>3+</sup> -bound | 51.8 | 6.4 | 41.7 |
| 19369 | Apo-form | 65.3 | 2.5 | 32.1 |
| 4581 | Apo-form | 59.5 | 3.6 | 36.9 |

The CB model prediction for the Ca^2+^ -bound form resulted in 56.9%, 6.5%, and 36.6% for the *α*-helix and *β*-sheet, and random coil, respectively. Venyaminov et al. [30] investigated the changes in the structure and stability of calbindin after calcium binding. They found a secondary structure content of 52% helix, 9% sheet, and 39% random coil using CD spectroscopy. Here, the CB model shows a maximum difference of 5% compared to results obtained via CD spectroscopy. Furthermore, the CB model predicts that calbindin-D28K, when bound to La^3+^, has a similar secondary structure, again consistent with current research that predicts that Ca^2+^ and La^3+^ lead to a similar protein structure [17]. Compared to the Ca^2+^ -bound form, the CB model predicts a 5% lower *α*-helix content for the La^3+^ -bound form. As the RMSE of the test dataset was 11% for *α*-helix, this slight underestimation is within the expected margin of error. For the linked PDB entry DSSP calculates 56% *α*-helix, 5.3% *β*-sheet, and 38.7% random coil content for La^3+^ -bound calbindin. This time, the model slightly underestimates the *α*-helix content by 4% and overestimates the random coil content by 3%. The predicted calbindin apo form tends to have more *α*-helix content than the Ca^2+^ - or La^3+^ -bound calbindin form. The two apo forms show 65.3% and 59.5% for *α*-helix, less *β*-sheet content with 2.5% and 3.6%, respectively. The DSSP calculation for the latter results in a *α*-helix content of 64.0%, 8.0% *β*-sheet and 28.0% random coil content. The increase in *α*-helix content is consistent with the DSSP calculations, but the decrease in *β*-sheet compared to the Ca^2+^ -bound form is not. The highest *α*-helix content was predicted for Mg^2+^ -bound calbindin at 69.6% and -0.4% for *β*-sheet. A prediction of -0.4% is obviously an incorrectly predicted percentage that represents a predicted nonexistence of the *β*-sheet. However, a negative prediction also affects the prediction of the other two secondary structure elements since the CB model has learned to predict an overall content of 100%. Therefore, either the *α*-helix content or the random coil content is overestimated by at least 0.4%. Research on the content of Mg^2+^ -bound calbindin secondary structures shows little information on secondary structure, but a study conducted by Skelton et al. [27] claimed that Mg^2+^ -as well as apo-forms have very similar secondary structure content to the Ca^2+^ -bound calbindin form.

### α-Synuclein

*α*-synuclein is an unfolded protein that accumulates in Lewy bodies and Lewy neurites in Parkinson’s and Alzheimer’s disease.[31] Depending on its environment and experimental condition, *α*-synuclein can undergo substantial conformational changes from high *β*-sheet content over extended *α*-helix regions to a mainly disordered structure. In figure 3 we investigated the structural changes of *α*-synuclein depending on the temperature.

McNulty et al. [18] detected a noticeable change in the ^15^N-HSQC spectra of *α*-synuclein when the temperature increased from 10^*°*^C to 30^*°*^C. Their investigation led them to suggest that the increase in temperature increases the content of ordered secondary structures (helix/sheet), leading to a change in the ^15^N-HSQC spectra. They assumed that the formation of the *α*-helix is the main factor for the increase of the secondary structure content. Our model, similar to the assumption of McNulty et al. [18], suggests an increase in *α*-helix formation with increasing temperature from 8.7% at 5^*°*^C to 22.1% at 30^*°*^C. But unlike the paper, the amount of secondary structure (which includes only the *α*-helix and the *β*-sheet) does not change significantly with the increase in temperature, as the *β*-sheet content decreases in the CB prediction.

In addition to the dynamic structural changes, the test cases also included five *α*-synuclein structures stated to have different secondary structures (Table 2).

**Table 2.**
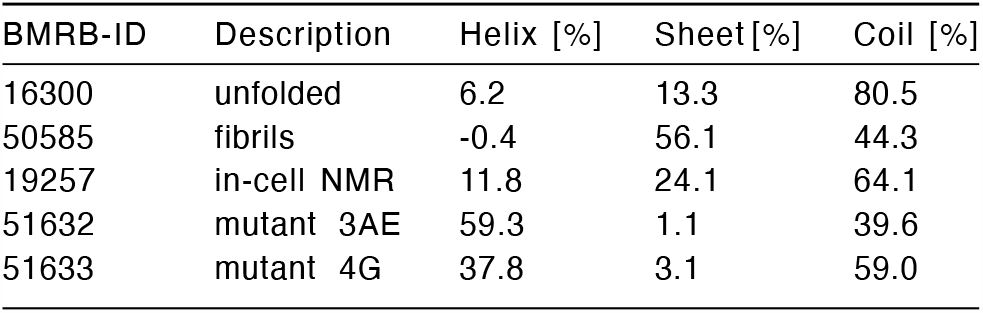
Secondary structure content prediction for different *α*-synuclein states. The description is the shortened version of the author’s BMRB protein description. The nomenclature of 3AE and 4G refers to a mutation of the residues 39 to 42 in the linker region from YVGS to either AAAE (3 AE mutant) or GGGG (4 G mutant).

For each of the different folds our CB model was able to identify the most abundant secondary structure element. For an unfolded *α*-synuclein (BMRB ID: 16300), a random coil fraction of 80.5% was predicted. Fibrils of *α*-synuclein fibril (in the presence of anionic phospholipids, BMRB ID: 50585) are predicted with our model to have a *β*-sheet fraction of 56.1% . These predictions are consistent with the high *β*-sheet content predicted by TALOS+ [3]. Two *α*-synuclein mutants with high *α*-helix content were also predicted. Although the authors did not provide an exact secondary structure content, their study postulated that mutant 3AE has a higher *α*-helix content than mutant 4G [8]. Our model predicts 59.3% for the *α*-helix content of the 3AE mutant and 37.8% for the *α*-helix content of the 4G mutant, consistent with the authors’ study results. Furthermore, for an in-cell measurement (BMRB ID: 19257), the CB model predicted a *α*-helix content of 11.8%, a *β*-sheet content of 24.1%, and a random coil content of 64.1%. Based on their measurements inside E. coli cells, Waudby et al. [32] concluded that the bacterial cytosol has the same properties as the disordered monomeric form observed in aqueous solution. Their NMR data were obtained at 277 K, comparable to the prediction for the 5^*°*^C measurement in temperature-dependent protein folding. The prediction based on the in-cell NMR spectrum of *α*-synuclein is 8.7% for *α*-helix, 23.5% for *β*-sheet, and 67.8% for random coil. With a maximum difference of 4%, the results agree and would thus support the claim that there is no structural change in the secondary structure of *α*-synuclein in the bacterial cytosol compared to its monomeric form.

### Summary

Overall, our model predicted changes in the secondary structure content induced by ligands or temperature with reasonable deviation compared to values that were determined with more complex, more expensive and/or more time-consuming methods. With more extensive BMRB/PDB datasets in the future, we think that this model and especially this approach of using unassigned N-HSQC spectra for secondary structure content prediction has big potential to be applied in in-cell NMR or to study ligand-induced conformational changes and protein-protein interactions.

## Acknowledgements

We thank Christopher Waudby (UCL School of Pharmacy) for discussing in-cell NMR applicability and Christoph Steinbeck (FSU Jena) for the awesome support. Parts of this work are financially supported by the Novartis Foundation for Medical-Biological Research (PB, Grant #22C222).

## Confilict of interest

None declared.

## Data availability

The spectral data and structure information utilized in this study can be accessed from the Biological Magnetic Resonance Bank (BMRB) and Protein Data Bank (PDB) databases. The web service is accessible via https://hsqc2struc.bellstedt-lab. ch. The CatBoost model as well as the TopSpin integration are available on GitHub (https://github.com/bellstedt-lab/hsqc2struc).

