## Supplementary Material for "HSQC2STRUC: A Machine Learning Model for Protein Secondary Structure Prediction using Unassigned NMR Spectra"

### Supplementary Information to "HSQC2STRUC: A Maschine Learning Model for Protein Secondary Structure Prediction using Unassinged NMR spectra"

Jonas Dietrich[1], and Peter Bellstedt[2]

[1]:Institute for Inorganic and Analytical Chemistry, Friedrich Schiller University (FSU), Lessinstrasse 8, 07743 Jena, Germany
[2]:University of Zurich (UZH) & University Hospital Zurich (USZ), Institute of Clinical Chemistry, Rämistrasse 100, 8091 Zurich, Switzerland

**Suppl. Figures** ...................Pages 1 to 9
**Suppl. Tables** ...................Pages 10 to 12
**Suppl. Listings**...................Page 13

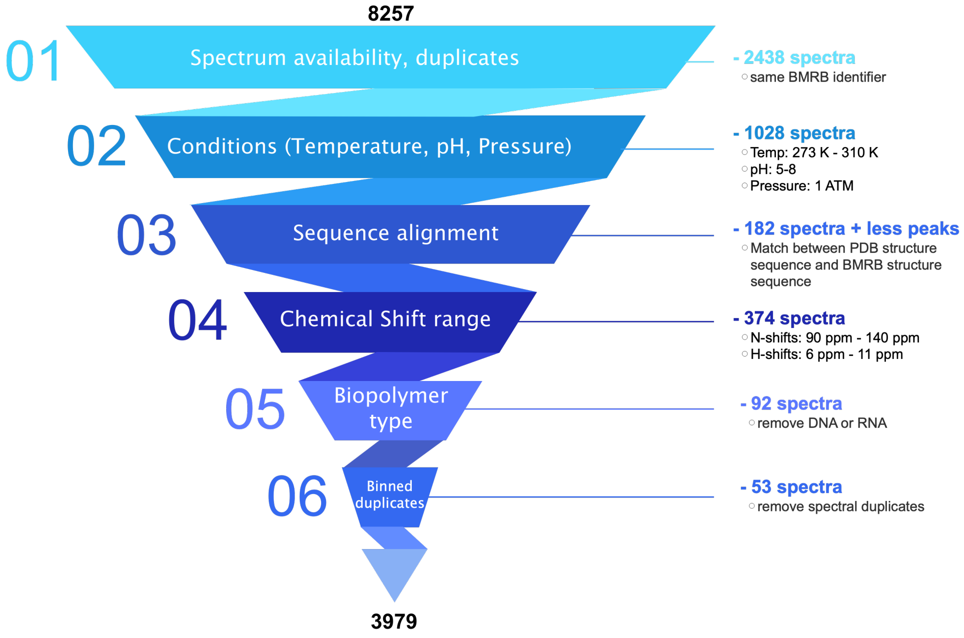


**Suppl. Figure 1**: Overview of the filtering process applied to the proteins from the BMRB-PDB match list. 3581 proteins were used for cross-validation and 398 proteins as the final test data set.

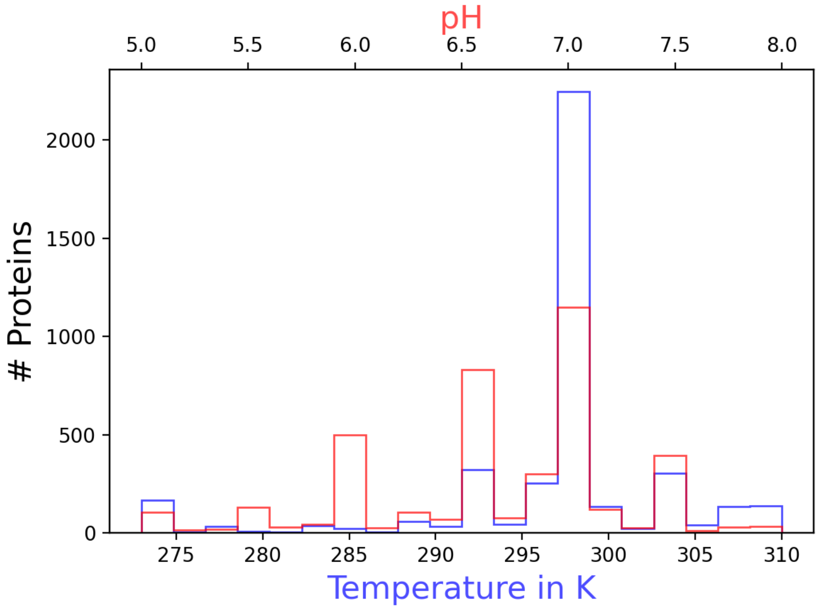


**Suppl. Figure 2**: Distribution of temperature (in blue) and sample pH (in red) of the proteins contained in the filtered data set.

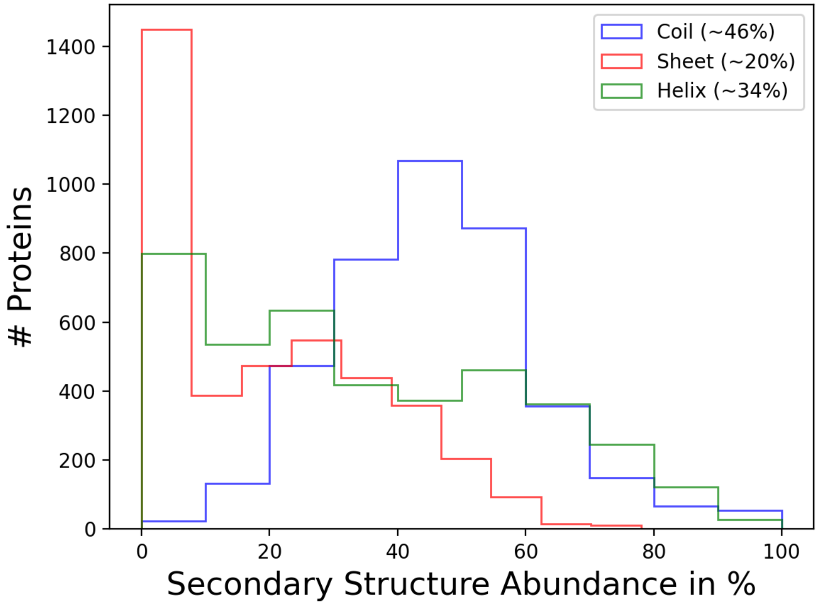


**Suppl. Figure 3**: Distribution of secondary structure content among the 3979 proteins used as input.

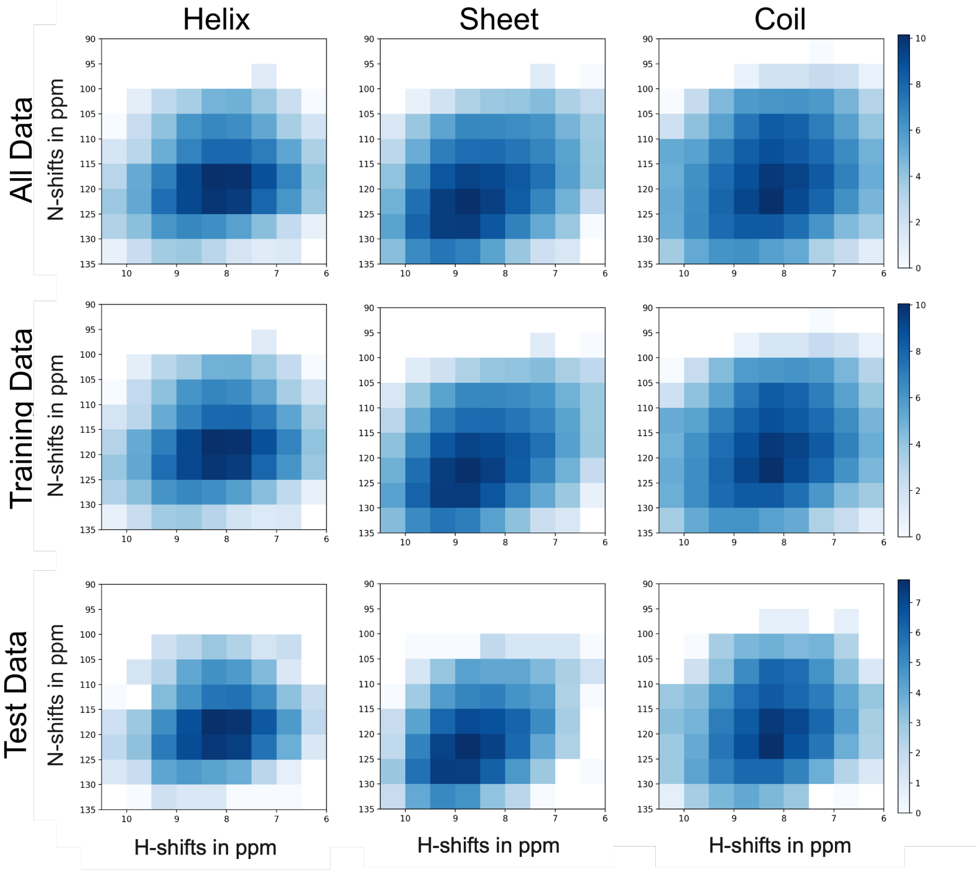


**Suppl. Figure 4**: Distribution of $^{1}$H, $^{15}$N-HSQC peaks belonging to the three different secondary structure elements and the different data sets. The overall density pattern of the training and test dataset is similar and therefore unbiased.

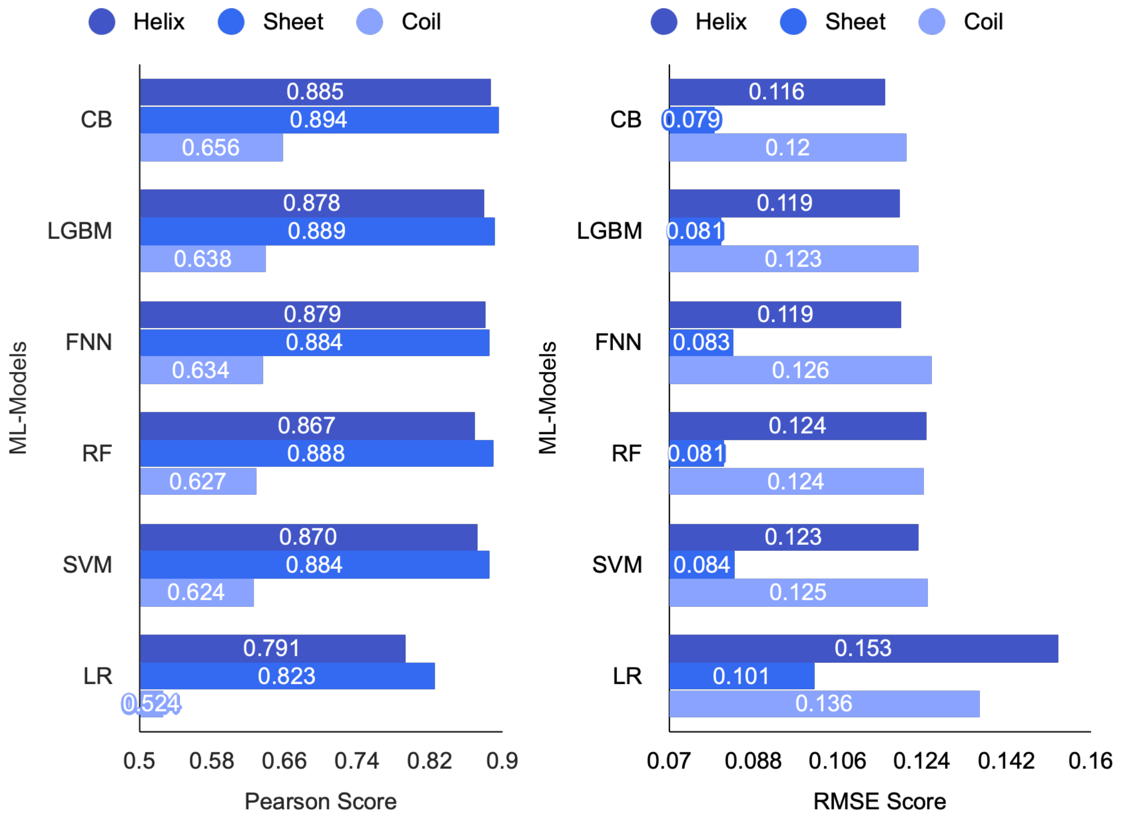


**Suppl. Figure 5**: Comparison of the Pearson values (left side) and Root-Mean-Square-Errors (RMSE, right side) for the prediction of secondary structure content of the proteins in the test data set and using the 5 (out of 13) best-performing machine learning models and linear regression (LR) as baseline model. CatBoost (CB) performed best (highest Pearson score and the lowest RMSE). LGBM: Light gradient boost machine, FNN: Feed forward neural network, RF: Random forest, SVM: Support vector machine, LR: Linear Regression.

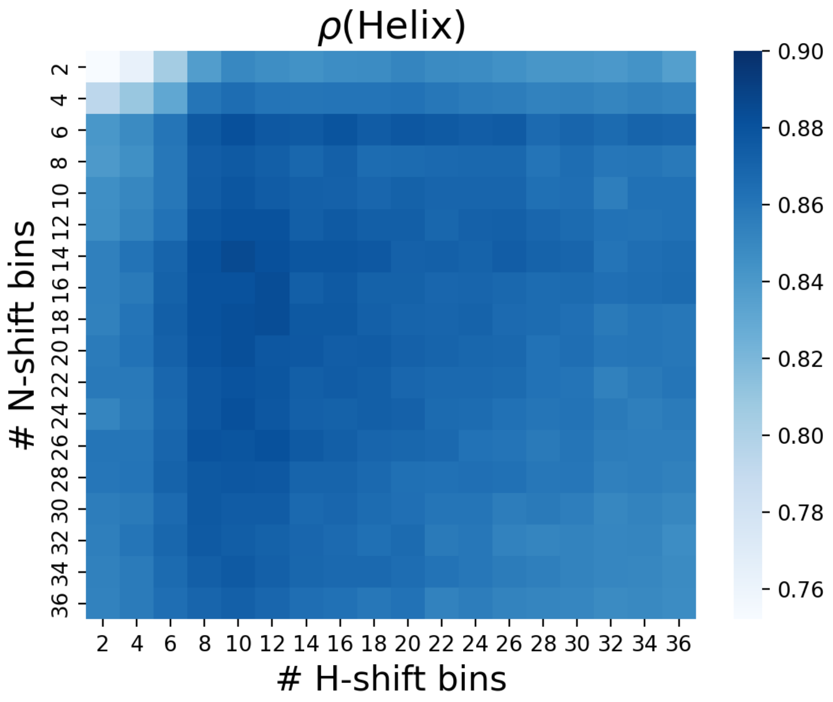


**Suppl. Figure 6**: Pearson scores of the CatBoost model using different binning sizes for the prediction of $\alpha$-helical content based on the 10-fold cross-validation with the training data set.

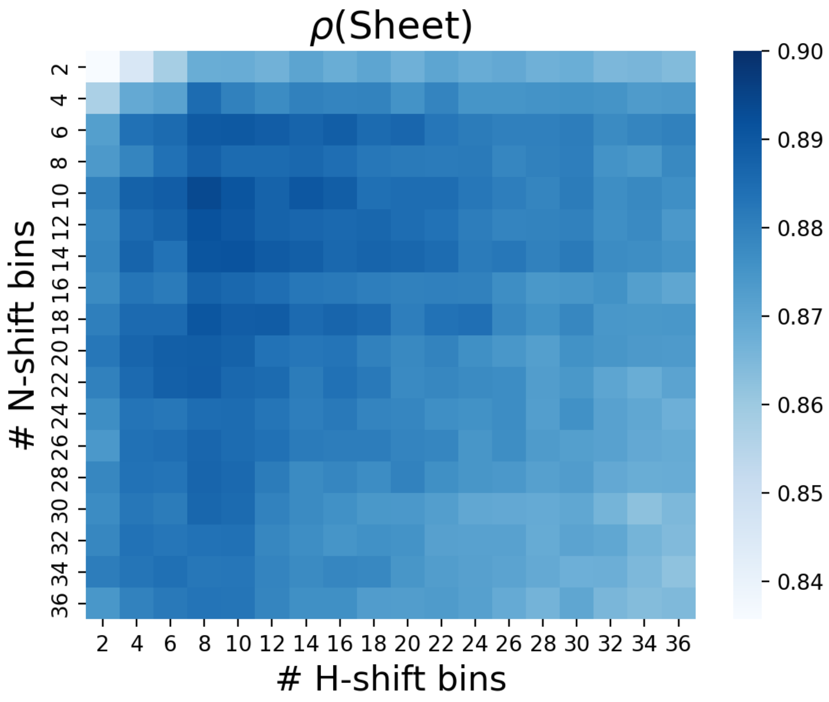


**Suppl. Figure 7**: Pearson scores of the CatBoost model using different binning sizes for the prediction of $\beta$-sheet content based on the 10-fold cross-validation with the training data set.

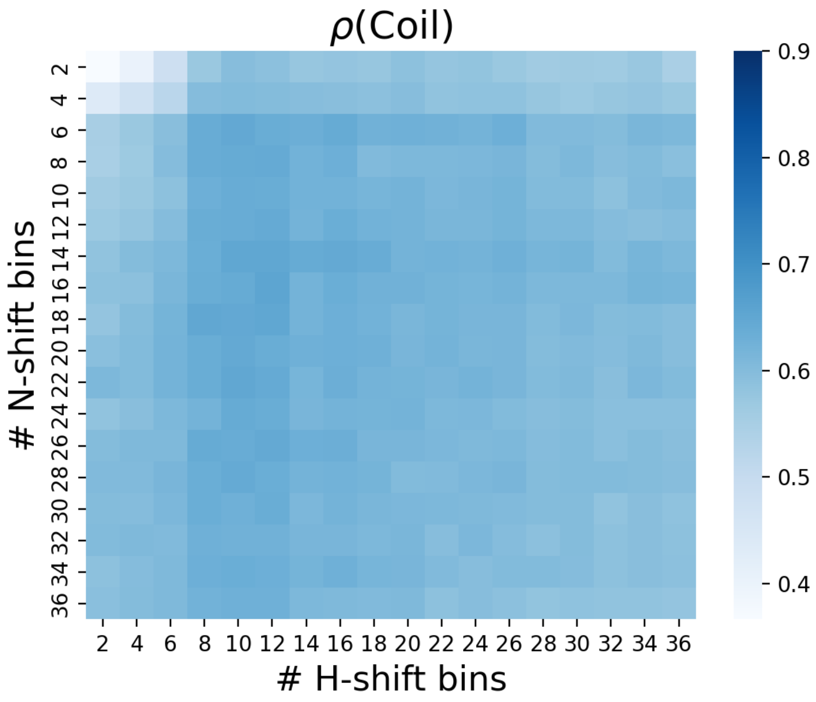


**Suppl. Figure 8**: Pearson scores of the CatBoost model using different binning sizes for the prediction of random coil content based on the 10-fold cross-validation with the training data set.

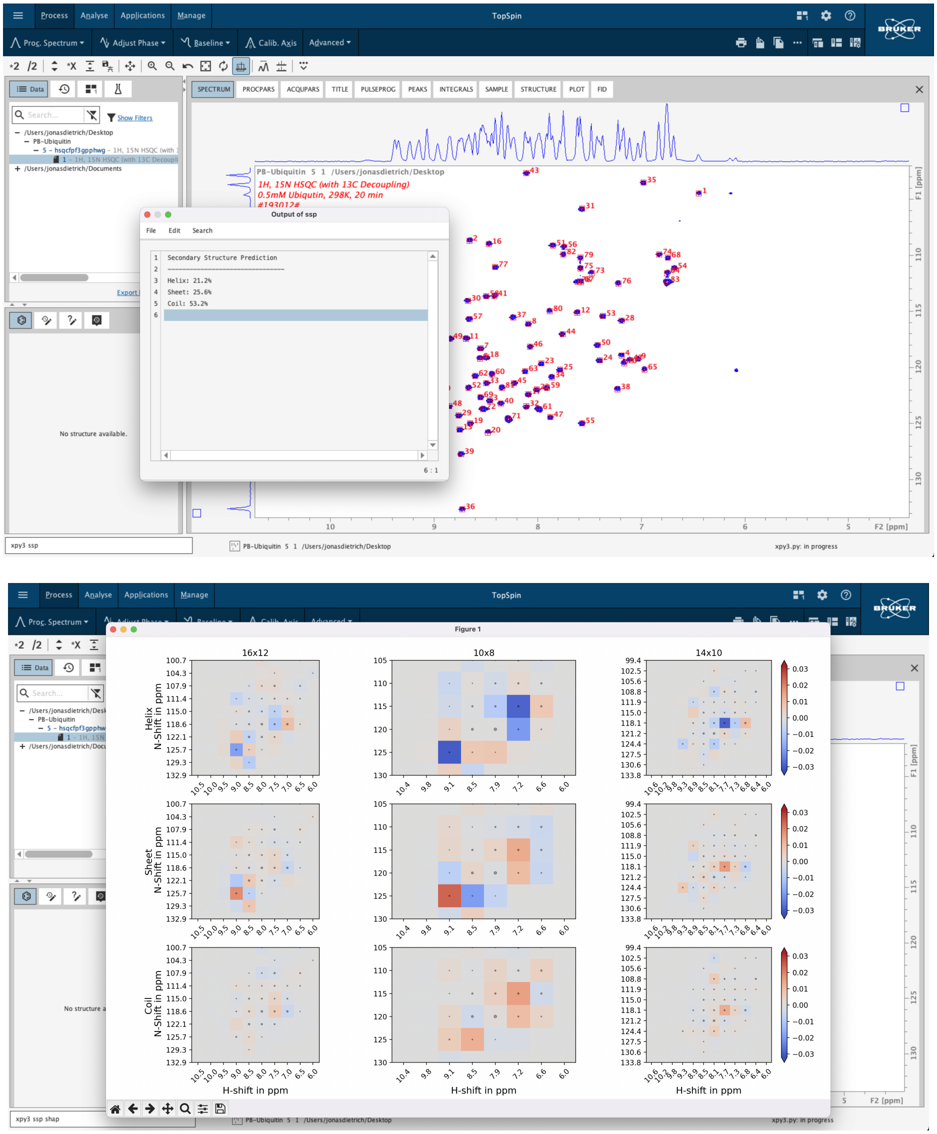


**Suppl. Figure 9**: Integration of our secondary structure prediction model in Bruker TopSpin (version 4.0 and above). After the scripts have been downloaded from our GitHub repository and after the necessary libraries are installed, the extension can be called via "xpy3 ssp". Please note, that peaks have to be picked before (e.g. via the pp command). After a few seconds, the prediction appears in a pop-up window (top image). To include SHAP values in the output (bottom image), use "xpy3 ssp shap" to start the prediction.

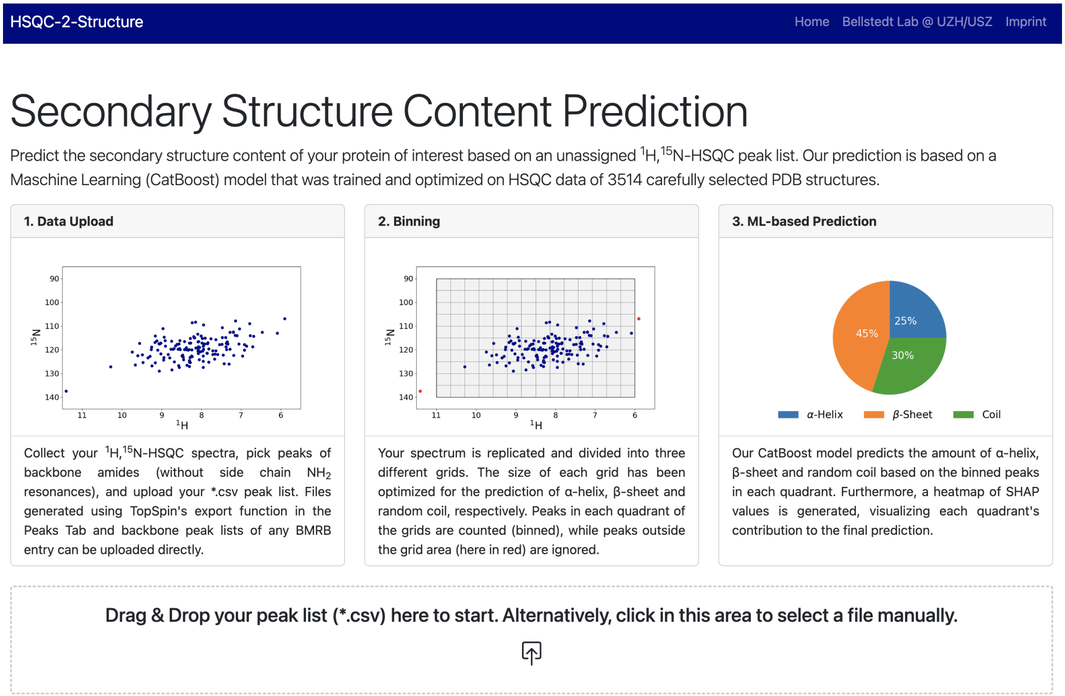


**Suppl. Figure 10**: Interface of the web service for predicting the secondary structure content. The underlying Python scripts are handled by Flask as a backend. The frontend design relies on the Bootstrap Framework. An Nginx web server is used as a proxy to handle the request efficiently. The web service is managed and run by a docker instance, which can be accessed via `https://hsqc2struc.bellstedt-lab.ch`.

**Suppl. Table 1.** Pearson and RMSE scores for $\alpha$-helix prediction for all 13 machine learning models tested, based on tenfold cross-validation on the training data set. For each model, only the result with the best-performing binning size is shown.

| Size | Model | Pearson | RMSE |
|---|---|---|---|
| 14x10 | CatBoost | 0.886 | 0.116 |
| 14x10 | LGBM | 0.879 | 0.119 |
| 34x34 | FeedForwardNeuralNetwork | 0.880 | 0.119 |
| 14x12 | SupportVectorMachine | 0.870 | 0.123 |
| 14x8 | BayesianNeuralNetwork | 0.873 | 0.123 |
| 6x12 | RandomForest | 0.867 | 0.124 |
| 6x10 | GradientBoost | 0.865 | 0.125 |
| 14x10 | XGB | 0.865 | 0.125 |
| 6x8 | KNN | 0.842 | 0.135 |
| 28x10 | RidgeRegression | 0.800 | 0.150 |
| 14x8 | LinearRegression | 0.791 | 0.153 |
| 6x12 | DecisionTree | 0.746 | 0.177 |
| 2x10 | ElasticNet | 0.712 | 0.186 |

**Suppl. Table 2.** Pearson and RMSE scores for $\beta$-sheet prediction for all 13 machine learning models tested, based on tenfold cross-validation on the training data set. For each model, only the result with the best-performing binning size is shown.

| Size | Model | Pearson | RMSE |
|---|---|---|---|
| 10x8 | CatBoost | 0.894 | 0.079 |
| 10x8 | LGBM | 0.889 | 0.081 |
| 6x8 | RandomForest | 0.888 | 0.081 |
| 10x8 | FeedForwardNeuralNetwork | 0.884 | 0.083 |
| 10x8 | BayesianNeuralNetwork | 0.882 | 0.085 |
| 14x10 | SupportVectorMachine | 0.882 | 0.084 |
| 6x12 | GradientBoost | 0.881 | 0.083 |
| 6x8 | XGB | 0.876 | 0.085 |
| 6x4 | KNN | 0.864 | 0.089 |
| 34x8 | RidgeRegression | 0.829 | 0.099 |
| 10x8 | LinearRegression | 0.823 | 0.101 |
| 2x2 | ElasticNet | 0.787 | 0.118 |
| 6x4 | DecisionTree | 0.779 | 0.118 |

**Suppl. Table 3.** Pearson and RMSE scores for random coil prediction for all 13 machine learning models tested, based on 10-fold cross-validation on the training data set. For each model, only the result with the best-performing binning size is shown.

| Size | Model | Pearson | RMSE |
|---|---|---|---|
| 16x12 | CatBoost | 0.656 | 0.120 |
| 14x10 | LGBM | 0.638 | 0.123 |
| 14x16 | FeedForwardNeuralNetwork | 0.634 | 0.126 |
| 6x8 | RandomForest | 0.627 | 0.124 |
| 14x10 | SupportVectorMachine | 0.623 | 0.125 |
| 14x8 | BayesianNeuralNetwork | 0.619 | 0.127 |
| 18x8 | GradientBoost | 0.605 | 0.127 |
| 18x8 | XGB | 0.604 | 0.128 |
| 14x10 | KNN | 0.562 | 0.133 |
| 30x16 | RidgeRegression | 0.551 | 0.133 |
| 14x8 | LinearRegression | 0.524 | 0.136 |
| 6x14 | DecisionTree | 0.392 | - |
| 6x10 | ElasticNet | 0.391 | 0.152 |

**Suppl. Table 4.** Best-performing binning sizes for different ML models and different secondary structure element predictions

| Model | Helix | Sheet | Coil |
|---|---|---|---|
| CB | 14x10 | 10x8 | 16x12 |
| LGBM | 14x10 | 10x8 | 14x10 |
| FNN | 34x34 | 10x8 | 14x16 |
| RF | 6x12 | 6x8 | 6x8 |
| SVM | 14x12 | 14x10 | 14x10 |
| LR | 14x8 | 10x8 | 14x8 |

**Suppl. Table 5.** Hyperparameter settings and optimization results. The column headings correspond to the names of the hyperparameters in Optuna without underscores. The loss function and the eval metric were set to "MultiRMSE".

| parameter | learning rate | depth | l2 leaf reg | border count | bagging temperature |
|---|---|---|---|---|---|
| min value | 0.001 | 4 | 1 | 32 | 0 |
| max value | 0.1 | 11 | 8 | 256 | 10 |
| opt parameter | 0.0319 | 7 | 6.5893 | 93 | 0.3297 |

**Suppl. Table 6.** Pearson scores $\rho$ of three independent models and one single model using all three best binning sizes as one input

| Model(s) | $\rho$(Helix) | $\rho$(Sheet) | $\rho$(Coil) |
|---|---|---|---|
| Independent | 0.884 | 0.893 | 0.645 |
| Single | 0.889 | 0.893 | 0.665 |
| Binning Size | 14x10 | 10x8 | 16x12 |

```
params = {'learning_rate': 0.1, 'depth': 6,
      'loss_function': 'MultiRMSE', 'eval_metric': 'MultiRMSE'}
```

**Suppl. Listing 1:** Python code with CatBoost parameter for 10x cross-validation on the training data set.

```
def ANN(input_size):
"""standard keras model"""
    model = keras.Sequential([
    layers.Dense(units=input_size, activation="relu"),
    layers.Dense(units=516, activation="relu"),
    layers.Dense(units=516, activation="relu"),
    layers.Dense(units=516, activation="relu"),
    layers.Dense(units=516, activation="relu"),
    layers.Dense(units=3),
])

model.compile(
    loss=keras.losses.Huber(),
    optimizer=keras.optimizers.Adam(learning_rate=0.001),
    metrics=[keras.metrics.MeanSquaredError()])

return model
...
KerasRegressor(build_fn=ANN_, epochs=10, batch_size=64)
```

**Suppl. Listing 2:** Python Code for the Feed Forward Neural Network Architecture.

```
def BNN(input_size):
    """standard keras model"""
    global tfd
    tfd = tfp.distributions

    # Define the Bayesian linear regression model
    inputs = keras.Input(shape=(input_size,))
    hidden = layers.Dense(516, activation="relu")(inputs)
    hidden = layers.Dense(516, activation="relu")(hidden)
    hidden = layers.Dense(516, activation="relu")(hidden)
    hidden = layers.Dense(516, activation="relu")(hidden)
    outputs = layers.Dense(3)(hidden)

    # Compile the model
    model = keras.Model(inputs=inputs, outputs=outputs)
    model.compile(loss=nll, optimizer=keras.optimizers.Adam(learning_rate=0.001))

    return model

# Define the likelihood as a Normal distribution
def likelihood(outputs):
     return tfd.Normal(loc=outputs, scale=1)

# Define the negative log likelihood loss function
def nll(y_true, y_pred):
    return -tf.reduce_mean(likelihood(y_pred).log_prob(y_true))
...

 KerasRegressor(build_fn=BNN_, epochs=10, batch_size=64)
```

**Suppl. Listing 3:** Python Code for Bayesian Neural Network Architecture.
